# MAB-5/Hox regulates the Q neuroblast transcriptome, including *cwn-1/Wnt,* to mediate posterior migration in *Caenorhabditis elegans*

**DOI:** 10.1101/2023.11.09.566461

**Authors:** Vitoria K. Paolillo, Matthew E. Ochs, Erik A. Lundquist

## Abstract

Neurogenesis involves the precisely-coordinated action of genetic programs controlling large-scale neuronal fate specification down to terminal events of neuronal differentiation. The Q neuroblasts in *C. elegans*, QL on the left and QR on the right, divide, differentiate, and migrate in a similar pattern to produce three neurons each. However, QL on the left migrates posteriorly, and QR on the right migrates anteriorly. The MAB-5/Hox transcription factor is necessary and sufficient for posterior Q lineage migration, and is normally expressed only in the QL lineage. To define genes controlled by MAB-5 in the Q cells, fluorescence-activated cell sorting was utilized to isolate populations of Q cells at a time in early L1 larvae when MAB-5 first becomes active. Sorted Q cells from *wild-type, mab-5* loss-of-function (*lof*), and *mab-5* gain-of-function (*gof*) mutants were subject to RNA-seq and differential expression analysis. Genes enriched in Q cells included those involved in cell division, DNA replication, and DNA repair, consist with the neuroblast stem cell identity of the Q cells at this stage. Genes affected by *mab-5* included those involved in neurogenesis, neural development, and interaction with the extracellular matrix. *cwn-1,* which encodes a Wnt signaling molecule, showed a paired response to *mab-5* in the Q cells: *cwn-1* expression was reduced in *mab-5(lof)* and increased in *mab-5(gof)*, suggesting that MAB-5 is required for *cwn-1* expression in Q cells. MAB-5 is required to prevent anterior migration of the Q lineage while it transcriptionally reprograms the Q lineage for posterior migration. Functional genetic analysis revealed that CWN-1 is required downstream of MAB-5 to inhibit anterior migration of the QL lineage, likely in parallel to EGL-20/Wnt in a non-canonical Wnt pathway. In sum, work here describes a Q cell transcriptome, and a set of genes regulated by MAB-5 in the QL lineage. One of these genes, *cwn-1,* acts downstream of *mab-5* in QL migration, indicating that this gene set includes other genes utilized by MAB-5 to facilitate posterior neuroblast migration.

## Introduction

Neurogenesis involves a precisely coordinated series of developmental events from early neuronal specification to terminal neuron-type differentiation. This program requires neurons to migrate from their birthplace and to settle in specific destinations where they will integrate into the neural circuit. Improper neuronal and neural crest migration and their subsequent failure to connect to the neural circuit can result in neurological and neurodevelopmental disorders. In humans, neuronal migration defects can cause symptoms such as seizures, poor motor function, and impaired cognitive development (reviewed in (BUCHSBAUM AND CAPPELLO 2019)). The *Caenorhabditis elegans* Q cells, bilateral neuroblasts born in the posterior-lateral region of the animal that undergo left-right asymmetric migration, are an excellent model to study directed neuronal migration.

The two Q cells are similar to one another in their morphology, initial position, lineage, and pattern of cell division, yet QR on the right side of the animal protrudes and migrates anteriorly while QL on the left protrudes and migrates posteriorly (SULSTON AND HORVITZ 1977; CHALFIE AND SULSTON 1981; MIDDELKOOP AND KORSWAGEN 2014; JOSEPHSON *et al*. 2016a; RELLA *et al*. 2016). Thus, distinct gene regulatory programs are activated in QR and QL to prompt L/R-asymmetry and induce migration patterns in opposite directions. Initial asymmetric migration is Wnt-independent and is regulated by the receptor molecules UNC-40/DCC and PTP-3/LAR, both of which are active in QL but not QR, leading to posterior versus anterior migrations (SUNDARARAJAN AND LUNDQUIST 2012; SUNDARARAJAN *et al*. 2014). The second phase of migration occurs after the first Q cell division and relies on Wnt signaling (CHALFIE *et al*. 1983; KENYON 1986; SALSER AND KENYON 1992; HARRIS *et al*. 1996; WHANGBO AND KENYON 1999; KORSWAGEN *et al*. 2000; HERMAN 2003; EISENMANN 2005; JI *et al*. 2013; JOSEPHSON *et al*. 2016a). QL descendants encounter a posterior EGL-20/Wnt signal secreted from the muscle and epidermal cells in the tail region. EGL-20 activates a canonical Wnt signaling pathway to induce expression of the Hox gene *mab-5* in QL and QL descendants, but not in QR and QR descendants. MAB-5 is both necessary and sufficient for posterior Q cell descendant migration, as QL descendants migrate anteriorly in *mab-5* loss-of-function (*lof*) mutants and QR descendants migrate posteriorly in *mab-5* gain-of-function (*gof*) mutants (JOSEPHSON *et al*. 2016a). The two phases of Q cell migration are independent, as initial QL and QR asymmetric migrations are normal in *mab-5* mutants (JOSEPHSON *et al*. 2016a). Furthermore, EGL-20 has a bipartite function in posterior QL migration: it first inhibits anterior migration via a non-canonical Wnt pathway, and activates *mab-5* expression via a canonical Wnt pathway. MAB-5 then maintains inhibition of anterior migration, and reprograms QL.a to migrate posteriorly (JOSEPHSON *et al*. 2016a). Thus, anterior migration must be inhibited to allow time for transcriptional responses of MAB-5 to reprogram QL.a to migrate posteriorly. Without inhibiting anterior migration, QL will migrate anteriorly.

MAB-5 promotes posterior migration of the QL descendants by directly inhibiting the expression of the Hox factor *lin-39* (WANG *et al*. 2013). As MAB-5 is not expressed in the QR descendants, LIN-39 drives the expression of genes such as the transmembrane protein *mig-13*, which acts cell-autonomously to promote anterior migration of the QR descendants. Apart from regulating *lin-39* and *mig-13* expression, it is not yet clear how MAB-5 functions to regulate Q cell migration. In an attempt to uncover additional MAB-5 targets, whole-animal RNA-seq on wild-type and *mab-5* mutants was previously conducted (TAMAYO *et al*. 2013). However, this analysis revealed potential *mab-5* targets expressed in non-Q cell types acting non-autonomously. For example, *mab-5* affects *spon-1/F-spondin* expression in posterior body wall muscle cells and SPON-1 functions non-autonomously to regulate Q cell migration (JOSEPHSON *et al*. 2016b). No candidates for *mab-5* targets in the Q cells themselves were evident from this approach, likely due to Q cell expression being a small subset of overall *mab-5-*regulated expression in other tissues (*e.g.,* muscles).

In this work, Q cells were sorted using fluorescence activated cell sorting and were subject to RNA-seq. Transcriptomes of Q cells from wild-type, *mab-5*(*lof*) mutants, and *mab-5*(*gof*) mutants were defined to identify genes regulated by *mab-5* in the Q cells. These studies revealed a diverse group of genes involved in neuronal differentiation, migration, and function. *cwn-1*, which encodes a Wnt molecule, showed a paired transcriptional response to *mab-5* in the Q cells, with reduced expression in *mab-5(lof)* and increased expression in *mab-5(gof)*. Functional studies showed that *cwn-1* is required downstream of *mab-5* to inhibit anterior migration, likely in an autocrine manner, and is part of the transcriptional network utilized by MAB-5 to prevent anterior migration.

## Materials and Methods

### Nematode strains

*C. elegans* strains were maintained by standard techniques and under standard conditions at 20℃. The following mutations and genetic constructs were utilized: LGII; *ayIs9[Pegl-17::gfp], cwn-1(ok546, gk130933, gk411542).* LGIII; *mab-5(gk670), mab-5(e1239), mab-5(lq165) mab-5(e1751).* LGIV; *egl-20(lq42, mu39).* LGV; *lqIs97[Pscm::mCherry], lqIs58[Pgcy-32::cfp]*. LGX; *lqIs221[Pegl-17::mab-5::gfp]*.

*mab-5(lq165)* was isolated in a forward genetic screen using ethyl methane sulfonate (EMS) for mutants with PQR migration defects. *mab-5(lq165)* was a C to T transition at chromosome III position 7,778,393, resulting in a premature stop codon at glutamine 160 (CAA to TAA). The sequence of the altered C to T in *mab-5(lq165)* is bold and underlined in the following sequence of the region: CATTGCATTTGACTGAAAGA**<u>C</u>**AAgtaagtctcactaccaga.

### Preparation of dissociated L1-stage larval cells

Q cells were isolated by L1 larval dissociation and fluorescence-activated cell sorting (FACS) from the following strains:

*mab-5(+)*

> LE4122: *ayIs9[Pegl-17::gfp] II; lqIs97[Pscm::mCherry] V*

*mab-5(loss-of-function)*

> LE5447: *mab-5(gk670) III; ayIs9[Pegl-17::gfp] II; lqIs97[Pscm::mCherry] V*
>
> LE5486: *mab-5(e1239) III; ayIs9[Pegl-17::gfp] II; lqIs97[Pscm::mCherry] V*

*mab-5(gain-of-function)*

> LE5657 *mab-5(e1751) III; ayIs9[Pegl-17::gfp] II; lqIs58 lqIs97[Pscm::mCherry] V*
>
> LE5487: *lqIs221[Pegl-17::mab-5::gfp]; ayIs9[Pegl-17::gfp] II; lqIs97[Pscm::mCherry] V*

L1 larval synchronization and cell isolation were completed as previously described with minor changes (SPENCER *et al*. 2014; TAYLOR *et al*. 2021). Strains were initially grown on 15-20 150-mm 8P nutrient agar plates seeded with *E. coli* strain NA22 per experiment (VON STETINA *et al*. 2007). Gravid adults and larvae were washed from the plates using sterile water when they had nearly depleted the bacterial food source, and allowed to settle in 50 ml conical tubes at room temperature. Excess water was removed, and embryos were isolated from gravid adults by treatment with 15-20 mL of freshly made bleach solution (18.75 mL of ddH_2_O, 5 mL of Clorox bleach, 1.25 mL 10 N NaOH) for ∼7 minutes with continuous vortexing, until the larvae and adults were completely dissociated and the embryos released. A series of five washes using M9 buffer were completed to remove the bleach solution. Each wash involved centrifuging for 2.5 minutes at 1300 RCF and removal of the supernatant. Embryos were then separated from debris by floating embryos on 30% cold sucrose solution with centrifugation, followed by a sterile water wash, then a M9 buffer wash. Embryos were allowed to hatch for 16-18 hours overnight in 2x50 ml conical tubes, each with 25 ml of M9, on a rotating shaker at 20℃.

Synchronized L1 animals were collected by centrifugation for 2.5 minutes at 1300 RCF and the supernatant discarded. The L1s were moved to 2x1.5 mL microcentrifuge tubes and pelleted for 2 minutes at 16000 RCF. One additional wash with ddH_2_O was completed to remove the M9 buffer, and residual ddH_2_O was removed to leave a compact L1 pellet of ∼40-100 µl per tube. For an ∼40 µl L1 pellet, 400 µl of freshly thawed SDS-DTT solution (ZHANG *et al*. 2011; ZHANG AND KUHN 2013) was added, and the mixture was pipetted slowly for 2 minutes at room temperature to remove clumps. Immediately after SDS-DTT treatment, 700 µl of egg buffer solution (ZHANG *et al*. 2011; ZHANG AND KUHN 2013) was added to the reaction and pipetted until mixed. Worms were pelleted for 1 minute at 16000 RCF and washed quickly 5 times with 1 mL of egg buffer to remove the SDS-DTT treatment. The SDS-DTT treated worms were moved to a 50 mL conical tube with 2 mL of 15 mg/mL of Pronase (Sigma-Aldrich, St. Louis, MO) in egg buffer at room temperature. A 21-gauge BD PrecisionGlide hypodermic needle on a 3-mL syringe was used for dissociation, by forcefully pulling and expelling the L1 animals through the needle. At 10 minutes, a sample was placed under a fluorescent dissecting microscope to monitor the progress of dissociation. In most cases, another 1 mL of Pronase solution was added, and vigorous dissociation via the needle/syringe was continued for another 10-15 minutes, while monitoring the progression of the dissociation every few minutes. Digestion with Pronase was stopped at 25 minutes regardless of the percent dissociation. In our hands, we found that this technique would dissociate between 60-90% of the L1 animals, depending on the strain. Even at the lower percent dissociation, we were able to obtain a sufficient number of sorted cells via FACS.

The dissociated L1 suspension was moved to 3x1.5 mL microcentrifuge tubes and pelleted at 10000 RCF for 3 minutes at 4℃. The supernatant was removed with a pipette, and the pelleted cells were washed twice in 1 mL of cold egg buffer solution. After the final wash, the cells were resuspended in 1 mL of cold egg buffer and the non-dissociated worms and large debris were pelleted at 100 RCF for 2.5 minutes at 4℃. The cell-containing supernatant was filtered through the top of a 35-micron filter-top tube on ice. Another 1 mL of egg buffer was added to the samples in the microcentrifuge tubes, centrifuged slowly at 100 RCF for 2.5 minutes at 4℃, and the supernatant filtered through the top of the 35-micron filter-top tube. A sample was checked under a fluorescent dissecting microscope and, depending on the concentration of cells, additional egg buffer was added to dilute the cells. We generally obtained 6-8 mL of cell suspension for sorting.

### Cell Sorting and RNA isolation

Upon completion of L1 animal dissociation, filtered cells were kept on ice in egg buffer. DAPI (1 µg/mL) was added prior to sorting to mark damaged cells. Sorting experiments were performed on a BD FACSAria Fusion equipped with a 70-micron diameter nozzle. For each experiment, 50,000 live cells from the whole animal were first sorted as a control sample; this sample included a random population of non-fluorescent, GFP+, mCherry+, and GFP+/mCherry+ cells. This was followed by a separate sorting of 20,000-50,000 live double positive Q cells (GFP+/mCherry+). In the wild-type strain LE4112, we observed that the Q cells (GFP+/mCherry+) made up 0.6±0.1% (3 separate experiments) of the total number of live cells. This is consistent with the number of Q cells present in L1-staged worms (∼0.4-0.7% or 2-4 Q cells or Q first division descendants of ∼558 total L1 cells).

Both control and Q cell samples were separately sorted into 750 µl of TRIzol LS Reagent (Invitrogen, #10296010), vortexed until mixed, and placed at -80℃ overnight. RNA was isolated using the Direct-zol RNA MiniPrep Kit (Zymo Research, #R2050). RNA integrity and concentration were determined using a High Sensitivity RNA ScreenTape in an Agilent TapeStation 2200 (Agilent Technologies). Our samples yielded 5-100ng of total RNA after RNA purification. Three biological replicates of each of the five strains resulted in 30 RNA samples that were used to construct RNA-seq libraries (3 biological replicates x 2 sorting (sorted Q cells and all cells) x five strains = 30 RNA samples).

### RNA-seq data collection and analysis

For each of the 30 samples, high-quality cDNA was generated from the total RNA using the SMART-Seq v4 Ultra Low Input RNA Kit (Takara Bio, Cat. #634889), and RNA-seq libraries were generated using the SMARTer ThruPLEX DNA-Seq Kit (Takara Bio, Cat. # R400674). Each library was indexed using the DNA Unique Dual Index Kit (Takara Bio, Cat. # R400665) for multiplexed sequencing using the Illumina NextSeq550 platform at the Genome Sequencing Core at the University of Kansas. The final libraries were validated and quantified via Qubit and TapeStation assays. Paired-end, 150 base-pair sequence reads were generated. Preprocessing of FASTQ files was completed using fastp (0.23.2) (CHEN *et al*. 2018), which included adapter trimming, per-read cutting by quality score, global trimming, and filtering out bad reads. Reads were aligned to the *C. elegans* reference genome [release WBcel235, version WBPS14 (WS271)] using HISAT2 (version 2.2.1) (KIM *et al*. 2015). Between 21 million and 85 million paired reads for each sample mapped to the *C. elegans* transcriptome. FASTQ files for this project are available in the Sequence Read Archive, Project Number PRJNA916208.

### Differential expression analysis

Gene counts were determined using featureCounts (version 2.0.1) (LIAO *et al*. 2014), and differential expression analysis was completed using DESeq2 (version 2.11.40.7) (LOVE *et al*. 2014). Genes with a false discovery rate (FDR) of less than 0.05 and a log2 fold change of 0.58 or greater (1.5 actual fold change) were considered significant.

### Functional annotation

For gene ontology term analysis and functional clustering, the Database for Annotation, Visualization, and Discovery (DAVID) (HUANG DA *et al*. 2009) was used. The Wormbase ID for each gene was determined, and gene lists were analyzed using the Functional Annotation Clustering tool in DAVID.

### Whole genome sequencing of *mab-5(e1751)*

Genomic DNA was isolated from *mab-5(e1751)*, and Illumina paired end libraries were constructed and subject to 150-cycle paired end sequencing. Reads were aligned to the reference genome using BWA-MEM2 (LI AND DURBIN 2009; LI AND DURBIN 2010). Resulting BAM files were analyzed in the Integrated Genome Viewer (ROBINSON *et al*. 2011; THORVALDSDOTTIR *et al*. 2012). Average read depth was ∼120, with a read depth of ∼350 in the chromosome 3 duplicated region (see Figure 5). *mab-5(e1751)* FASTQ files can be found in the Sequence Read Archive, project number PRJNA1028545.

### Scoring AQR and PQR position

AQR and PQR migration defects were visualized and quantified using previously described techniques (CHAPMAN *et al*. 2008; SUNDARARAJAN AND LUNDQUIST 2012). The transgene *lqIs58[Pgcy-32::CFP]* was used to visualize AQR and PQR along the anterior-posterior (AP) axis of L4 or young adult animals. Five regions along the AP axis were used to score the final positions of AQR and PQR. Position 1 is located around the posterior pharyngeal bulb and is the wild-type location of AQR. Position 2 is a region posterior to position 1 but anterior to the vulva. Position 3 is a region centered around the vulva. Position 4 includes the region where the Q neuroblasts are born near the posterior deirid ganglion. Position 5 is located at the very posterior of the worm and is the final location of wild-type PQR, which settles just posterior to the anus. At least 100 animals of each genotype were quantified using the above scoring method. Fisher’s exact test was used to determine significance.

## Results

### Fluorescence activated cell sorting and transcriptome sequencing of Q cells

The bilateral Q neuroblasts, the sisters of the V5 hypodermal seam cells, are born in the posterior-lateral region of the animal, with QL on the left and QR on the right (Figure 1) (CHAPMAN *et al*. 2008; MIDDELKOOP AND KORSWAGEN 2014). QR extends a protrusion anteriorly over the V4 seam cell, and QL posteriorly over the V5 seam cell (Figure 1A). As QL extends posteriorly, it encounters an EGL-20/Wnt signal from posterior cells. Via canonical Wnt signaling, EGL-20/Wnt activates the expression of MAB-5/Hox in QL but not QR (Figure 1B and C) (CHALFIE *et al*. 1983; KENYON 1986; SALSER AND KENYON 1992; HARRIS *et al*. 1996; WHANGBO AND KENYON 1999; KORSWAGEN *et al*. 2000; HERMAN 2003; EISENMANN 2005; JI *et al*. 2013). The nuclei and cell bodies of the Q cells follow the protrusion and migrate on top of the V4 and V5 seam cells, at which time they undergo their first division (Figure 1B and C). MAB-5/Hox is required to prevent anterior migration of QL.a/p and drive further posterior migration of QL.a (JOSEPHSON *et al*. 2016a). In *mab-5*(*lof*) mutants, PVM, SDQL, and PQR migrate anteriorly (Figure 1D). Ectopic MAB-5/Hox expression in QR is sufficient to prevent anterior QR.a/p migration and to promote posterior QR.a migration (TAMAYO *et al*. 2013). In *mab-5*(*gof*) mutants, AVM, SDQR, and AQR migrate anteriorly (Figure 1E). Thus, MAB-5/Hox is a terminal selector determinant of posterior Q neuroblast migration.

**Figure 1:**
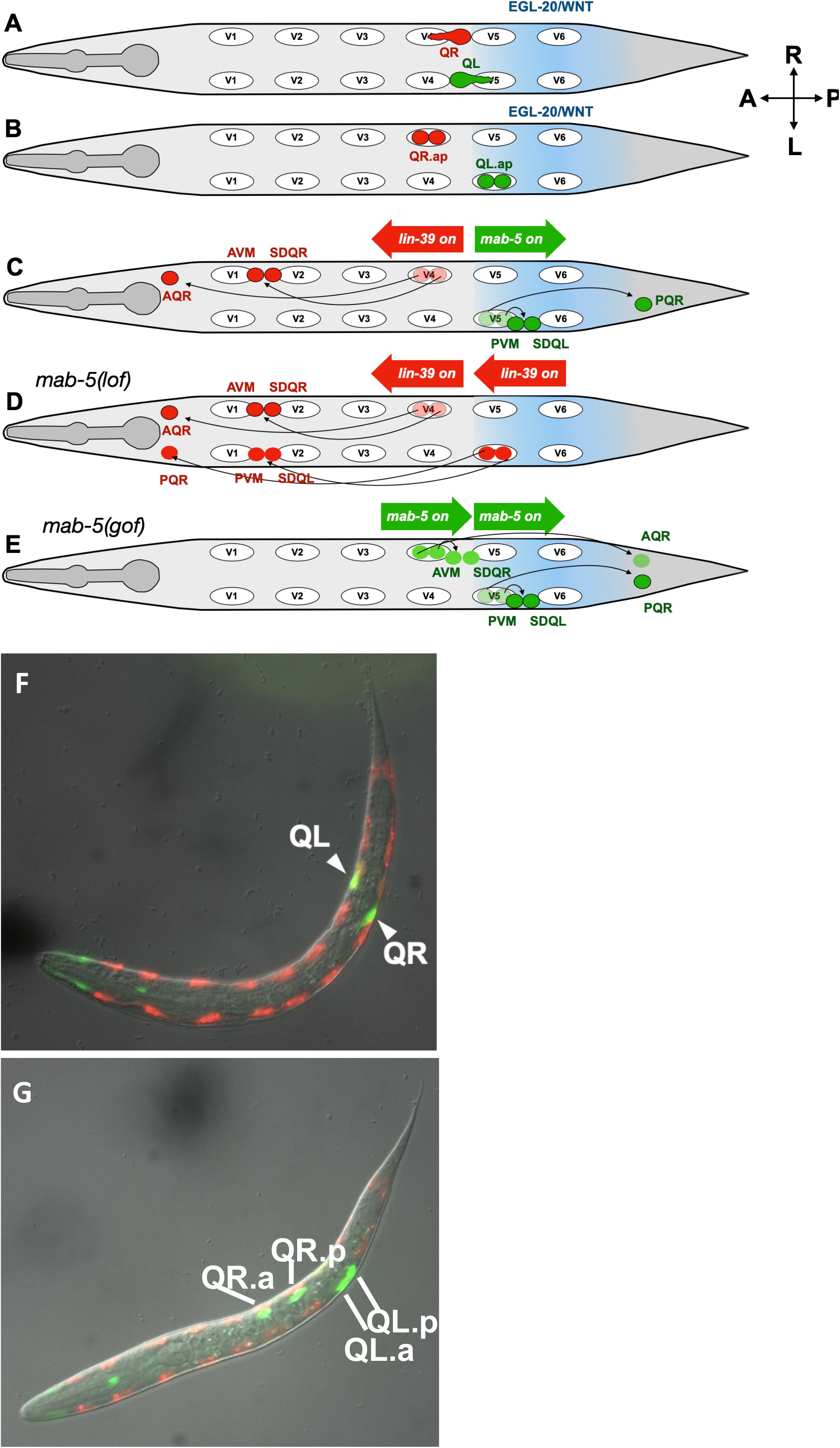
Migration of the Q neuroblasts and their descendants. A) Schematic of initial, Wnt-independent migration of QR and QL at 1.5-3h post-hatching. B) First cellular division of the Q neuroblasts at 5h post hatching to generate QR.ap and QL.ap. Wnt-dependent migration of the Q neuroblast descendants. EGL-20/WNT signaling activates the Hox gene *mab-5* in QL descendants, but not in QR descendants. MAB-5 function is necessary and sufficient for posterior migration. MAB-5 inhibits expression of the Hox gene *lin-39*; thus, *lin-39* is expressed in the absence of MAB-5 in the QR descendants and is required for anterior migration. D) In a *mab-5* loss-of-function mutant, *lin-39* is expressed in QL.a/p, and they migrate anteriorly. E) In a *mab-5* gain-of-function mutant, *mab-5* is expressed in QR.a/p, resulting in repression of *lin-39* and posterior migration. F) and G) Merged fluorescence and DIC micrographs of starved and arrested L1 wild-type animals. These animals express *Pscm::mCherry* in the seam cells and Q cells, and *Pegl-17::gfp* in the Q cells and some cells in the anterior. F) In this starved and arrested L1 lava, the Q cells have begun their initial migrations. G) In this starved and arrested L1 larva, the Q cells have complete initial migration and have undergone the first cell division. QR.a/p have started migrating to the anterior.

As no promoter is known to be expressed exclusively in the Q cells, overlapping *gfp* and *mCherry* expression driven by two different promoters was utilized to sort Q cells. *egl-17* is expressed in the Q cells, pharyngeal cells, and pharyngeal neurons (BURDINE *et al*. 1998), while the seam cell marker (*scm*) is expressed in the hypodermal seam cells (HOPE 1991; GENDREAU *et al*. 1994) and the Q cells (CHAPMAN *et al*. 2008) (Figure 1F and G). The Q cells are the only cells that express both *egl-17* and *scm*. For this reason, *ayIs9[Pegl-17::gfp]* and *lqIs97[Pscm::mCherry]* were included in all strains subject to FACS (Figure 2A and B and Materials and Methods).

**Figure 2.**
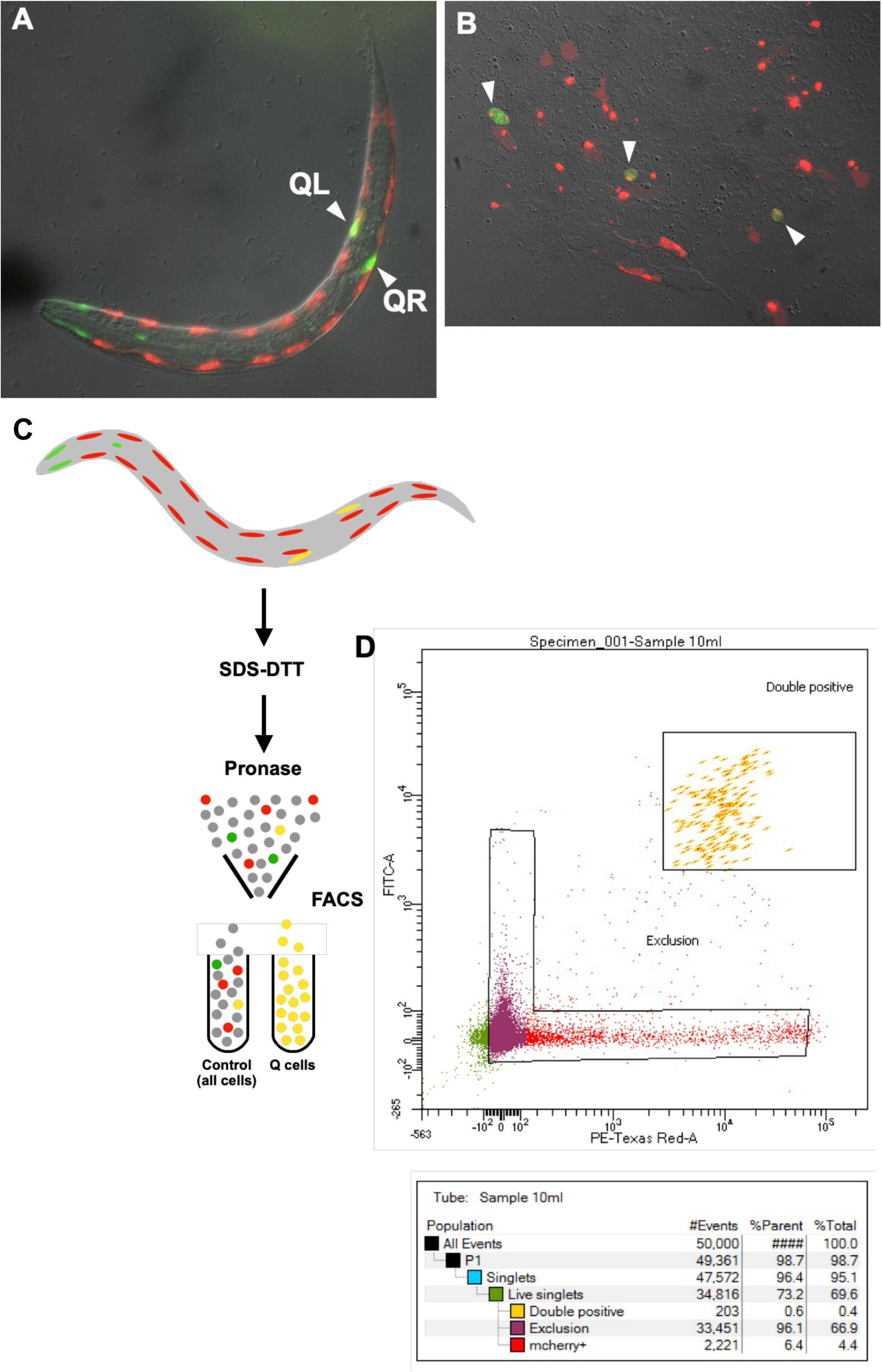
Larval dissociation and fluorescence-activated cell sorting of Q cells. A) A starved and arrested L1 larva expressing *Pscm::mCherry* and *Pegl-17::gfp* as described in Figure 1 (this is the identical image in Figure 1F). B) A field of cells after larval dissociation, with *Pscm::mCherry* and *Pegl-17::gfp* expression. *mCherry/gfp* double positive cells Q cells are indicated with arrowheads. C) A diagram of the SDS/DTT and Pronase larval dissociation and FACS sorting of double-positive Q cells. The results of a FACS gating sort to define the parameters of double-positive Q cells.

Q cells were isolated from starved and arrested L1-stage animals using a two-color sorting protocol based on (SPENCER *et al*. 2014; TAYLOR *et al*. 2021) (Figure 2 and Materials and Methods). In starved and arrested L1 larvae, all Q cells had begun their initial migration (Figure 1F), some had completed initial migration and undergone the first division, and QR.a and QR.p had begun migrating in some animals (Figure 1G). Thus, sorted cells included a mix of QL, QR, and the daughter cells QX.a and QX.p. *mab-5* is first expressed in QL as it undergoes its initial migration process of migration, so the effects of *mab-5* on the Q cells should be evident in these starved and arrested L1 animals. Indeed, *mab-5* expression was enriched in sorted Q cells (Supplemental File 1).

FACS was conducted on synchronized and dissociated L1 larva as described in Materials and Methods (SPENCER *et al*. 2014; TAYLOR *et al*. 2021). After L1 dissociation and filtering, 50,000 live cells were sorted to be used as an all-cell control, and 20,000-50,000 double-positive cells (Q cells) were sorted for each experiment. Three biological replicates of the sorting experiment per strain were completed (for both the Q cell and control all-cell populations).

RNA-seq libraries were sequenced using the Illumina NextSeq550 platform with paired-end, 150 base-pair sequencing (see Materials and Methods). Reads were aligned to the *C. elegans* reference genome using HISAT2 and counted using featureCounts (see Materials and Methods).

### The Q cell transcriptome

DEseq2 (see Materials and Methods) was used to compare wild-type Q cell expression to all cells. The wild-type Q cell transcriptome was identified by filtering out lowly expressed genes with a DEseq2 normalized base mean of less than 10 among all 46,748 gene elements, which includes protein coding genes and genes encoding RNAs. This resulted in 20,110 gene elements expressed in the Q cells (Supplemental File 1; WTQ x WTAll tab). Among these were genes known to be expressed and function in the Q cells, including *mab-5, lin-39, mig-13,* and *unc-40*.

### Differentially expressed genes in wild-type Q cells relative to the whole animal

DEseq2 was used to determine expression differences between sorted Q cells and the all-cells control from wild-type animals (*q*-value ≤ 0.05, log2-fold change ≥ 0.58, base mean expression ≥ 10). This analysis revealed that expression of 3,101 genes was significantly enriched in Q cells, and 2,560 significantly depleted (Supplemental File 1; Q cell enriched and Q cell depleted tabs). As expected, genes known to be expressed in QL, QR, QLap, and/or QRap, as determined by anatomy terms on WormBase, were significantly enriched in Q cells (Figure 3). This list included *mab-5* and *lin-39*, which control QL and QR migration, respectively.

**Figure 3.**
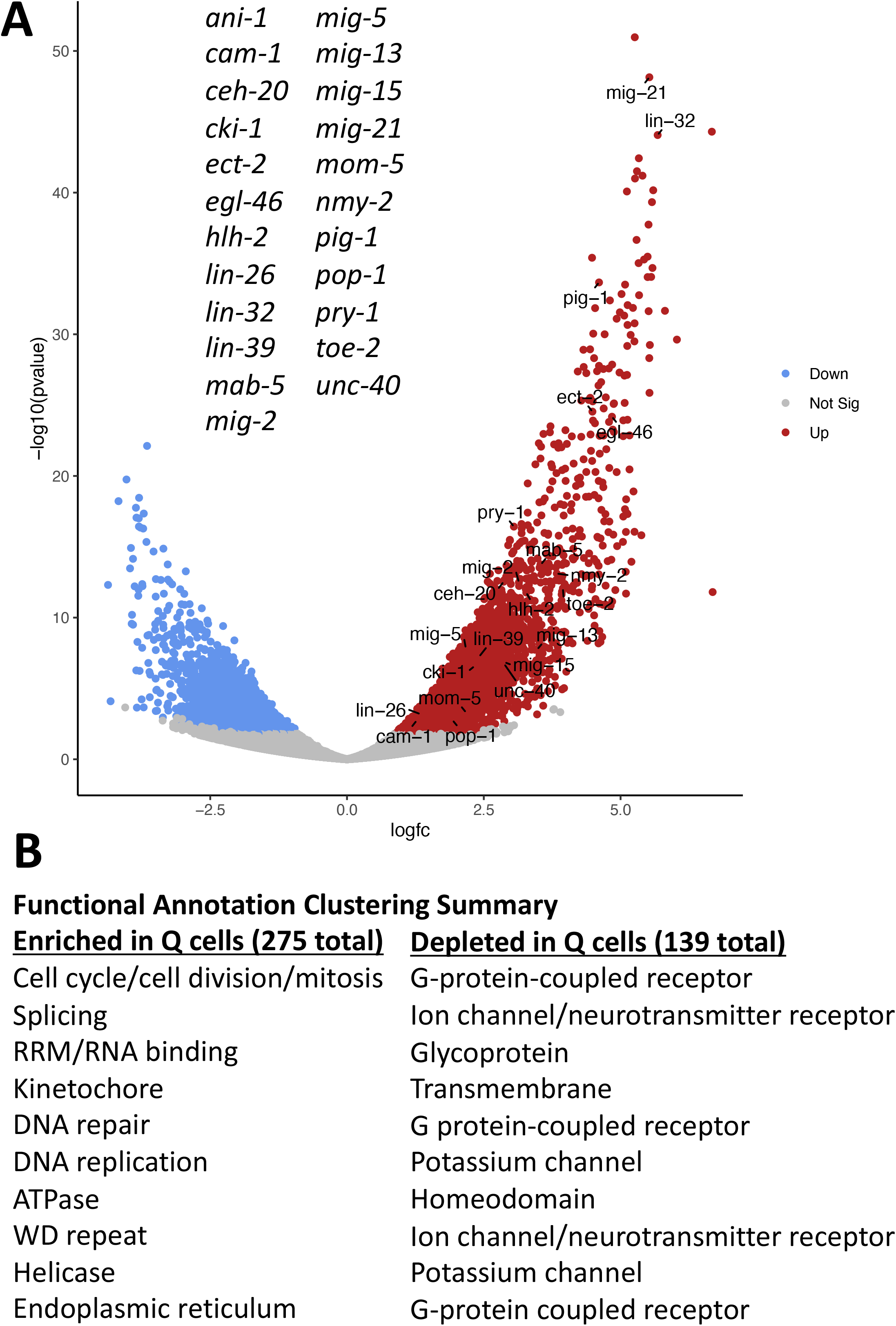
Genes differentially expressed in sorted Q neuroblasts. A) A volcano plot of genes differentially expressed in Q cells versus all cells is shown. The X axis is the log fold change in expression in Q cells compared to all cells. The Y axis is the -log10 *p* value significance of the expression change. Genes enriched in Q cells are red, and those depleted in Q cells are blue. Genes with known Q cell expression are listed and indicated (genes with Q cell anatomy terms in Wormbase). B) The top ten most significant functional clusters of genes enriched or depleted in Q cells (Supplemental File 3)

The Database for Annotation, Visualization, and Discovery (DAVID) (HUANG DA *et al*. 2009) was used categorize genes into significantly-enriched functional clusters using the Functional Annotation Cluster tool. Genes enriched in Q cells defined 275 functional clusters (Supplemental File 3). The ten most significant functional clusters are shown in Figure 3B, and include terms characteristic of cell division, DNA replication, and DNA repair, consistent with the mitotic stem cell nature of the Q cells at this time in development. Genes depleted in the Q cells clustered into 139 groups (Supplemental File 3). The ten most significant included terms involved in neuronal differentiation and function, such as G protein coupled receptors and neurotransmitter receptors. This is consistent with the Q cells at this time in development being in a mitotic stem cell state with neuronal differentiation possibly repressed.

### Differentially expressed genes in *mab-5* loss-of-function mutant Q cells

Q cells were sorted from two *mab-5* loss of function mutants, *gk670* and *e1239. mab-5(gk670)* is a deletion (TAMAYO *et al*. 2013), and *mab-5(e1239)* is a splicing mutation resulting in the use of a cryptic spice site, a frame shift, and introduction of a premature stop codon (Wormbase). Both result in complete failure of PQR posterior migration, with 99% in both mutants migrating anteriorly to the normal position of AQR (Table 1). *mab-5(lq165)* introduced a premature stop codon at Q160 in *mab-5* (see Materials and Methods) and had a similar phenotype (99% of PQR at position 1) (Table 1).

**Table 1.**
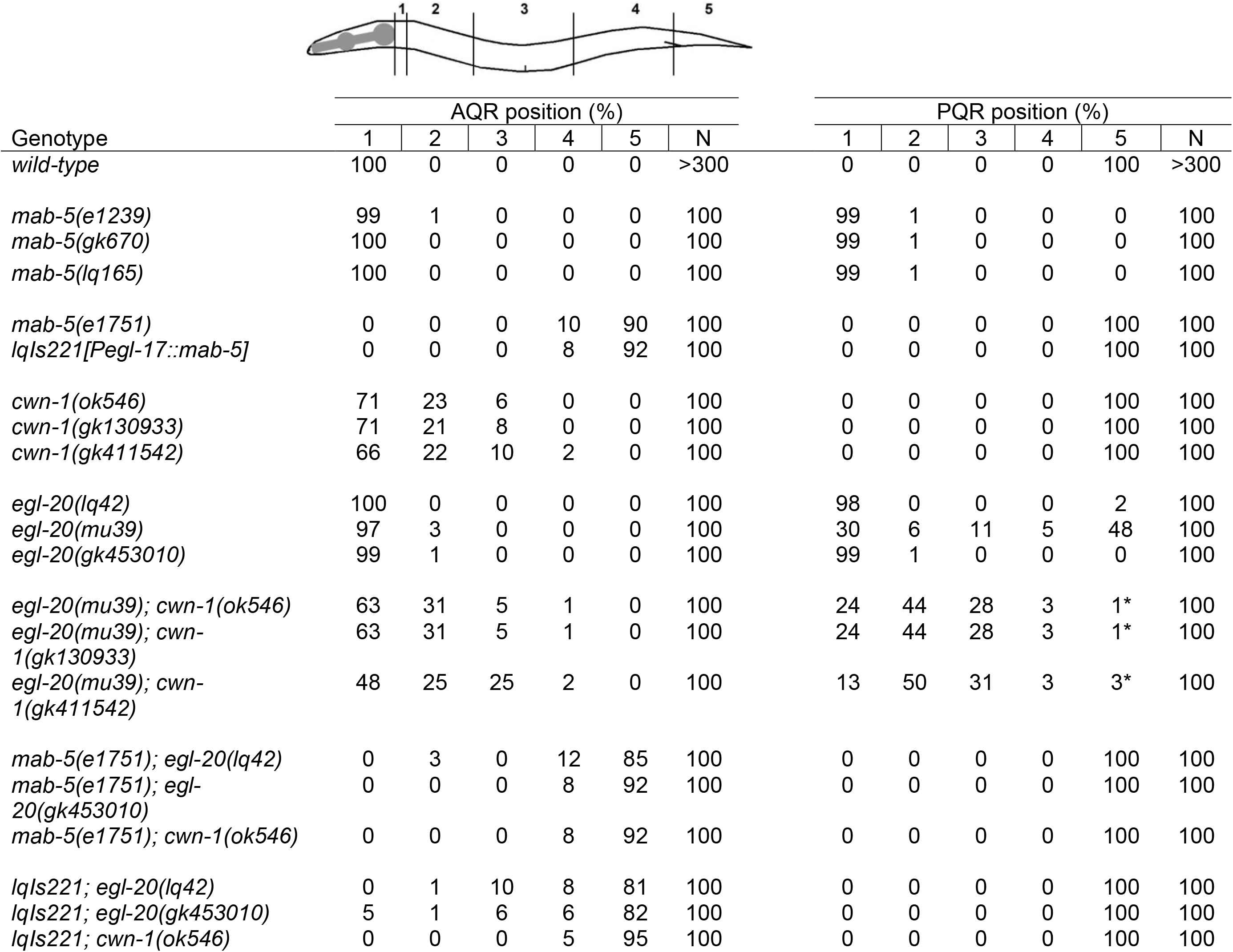
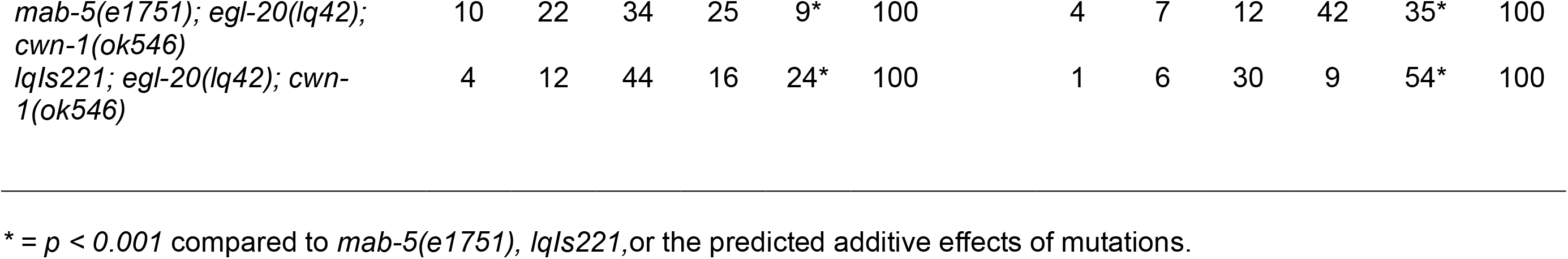
AQR and PQR migration.

| Genotype | AQR position (%) |  |  |  |  |  | PQR position (%) |  |  |  |  |  |
| --- | --- | --- | --- | --- | --- | --- | --- | --- | --- | --- | --- | --- |
|  | 1 | 2 | 3 | 4 | 5 | N | 1 | 2 | 3 | 4 | 5 | N |
| *wild-type* | 100 | 0 | 0 | 0 | 0 | >300 | 0 | 0 | 0 | 0 | 100 | >300 |
| *mab-5(e1239)* | 99 | 1 | 0 | 0 | 0 | 100 | 99 | 1 | 0 | 0 | 0 | 100 |
| *mab-5(gk670)* | 100 | 0 | 0 | 0 | 0 | 100 | 99 | 1 | 0 | 0 | 0 | 100 |
| *mab-5(lq165)* | 100 | 0 | 0 | 0 | 0 | 100 | 99 | 1 | 0 | 0 | 0 | 100 |
| *mab-5(e1751)* | 0 | 0 | 0 | 10 | 90 | 100 | 0 | 0 | 0 | 0 | 100 | 100 |
| *lqls221[Pegl-17::mab-5]* | 0 | 0 | 0 | 8 | 92 | 100 | 0 | 0 | 0 | 0 | 100 | 100 |
| *cwn-1(ok546)* | 71 | 23 | 6 | 0 | 0 | 100 | 0 | 0 | 0 | 0 | 100 | 100 |
| *cwn-1(gk130933)* | 71 | 21 | 8 | 0 | 0 | 100 | 0 | 0 | 0 | 0 | 100 | 100 |
| *cwn-1(gk411542)* | 66 | 22 | 10 | 2 | 0 | 100 | 0 | 0 | 0 | 0 | 100 | 100 |
| *egl-20(lq42)* | 100 | 0 | 0 | 0 | 0 | 100 | 98 | 0 | 0 | 0 | 2 | 100 |
| *egl-20(mu39)* | 97 | 3 | 0 | 0 | 0 | 100 | 30 | 6 | 11 | 5 | 48 | 100 |
| *egl-20(gk453010)* | 99 | 1 | 0 | 0 | 0 | 100 | 99 | 1 | 0 | 0 | 0 | 100 |
| *egl-20(mu39); cwn-1(ok546)* | 63 | 31 | 5 | 1 | 0 | 100 | 24 | 44 | 28 | 3 | 1* | 100 |
| *egl-20(mu39); cwn-1(gk130933)* | 63 | 31 | 5 | 1 | 0 | 100 | 24 | 44 | 28 | 3 | 1* | 100 |
| *egl-20(mu39); cwn-1(gk411542)* | 48 | 25 | 25 | 2 | 0 | 100 | 13 | 50 | 31 | 3 | 3* | 100 |
| *mab-5(e1751); egl-20(lq42)* | 0 | 3 | 0 | 12 | 85 | 100 | 0 | 0 | 0 | 0 | 100 | 100 |
| *mab-5(e1751); egl-20(gk453010)* | 0 | 0 | 0 | 8 | 92 | 100 | 0 | 0 | 0 | 0 | 100 | 100 |
| *mab-5(e1751); cwn-1(ok546)* | 0 | 0 | 0 | 8 | 92 | 100 | 0 | 0 | 0 | 0 | 100 | 100 |
| *lqls221; egl-20(lq42)* | 0 | 1 | 10 | 8 | 81 | 100 | 0 | 0 | 0 | 0 | 100 | 100 |
| *lqls221; egl-20(gk453010)* | 5 | 1 | 6 | 6 | 82 | 100 | 0 | 0 | 0 | 0 | 100 | 100 |
| *lqls221; cwn-1(ok546)* | 0 | 0 | 0 | 5 | 95 | 100 | 0 | 0 | 0 | 0 | 100 | 100 |

|  |  |  |  |  |  |  |  |  |  |  |  |  |  |
| --- | --- | --- | --- | --- | --- | --- | --- | --- | --- | --- | --- | --- | --- |
| *mab-5(e1751); egl-20(lq42);*<br>*cwn-1(ok546)* | 10 | 22 | 34 | 25 | 9* | 100 |  | 4 | 7 | 12 | 42 | 35* | 100 |
| *lqls221; egl-20(lq42); cwn-*<br>*1(ok546)* | 4 | 12 | 44 | 16 | 24* | 100 |  | 1 | 6 | 30 | 9 | 54* | 100 |

\* = $p < 0.001$ compared to *mab-5(e1751)*, *lqls221*, or the predicted additive effects of mutations.

DEseq2 was used to identify differentially expressed genes in *mab-5* mutant versus wild-type Q cells (*q*-value ≤ 0.05, log2-fold change ≥ 0.58, base mean expression ≥ 10) (Supplemental File 2; gk670_x_WT and e1239_x_WT tabs). In *mab-5(gk670),* 218 genes showed reduced expression compared to wild-type, and 72 showed increased expression (Supplemental File 2; gk670_x_WT_DOWN and gk670_x_WT_UP tabs). In *mab-5(e1239)*, 27 genes showed reduced expression and 50 genes increased expression (Supplemental File 2; e1239_x_WT_DOWN and e1239_x_WT_UP tabs). Seven genes showed reduced expression in both *mab-5(gk670)* and *mab-5(e1239)* (*M162.5, sem-2, ZK512.1, vab-8, E01G4.5, lon-1,* and *unc-75*), and five genes showed increased expression in both (*C01H6.8, T13H10.2, T25G12.3, nhr-127,* and *21ur-2841*) (Table 2).

**Table 2.**
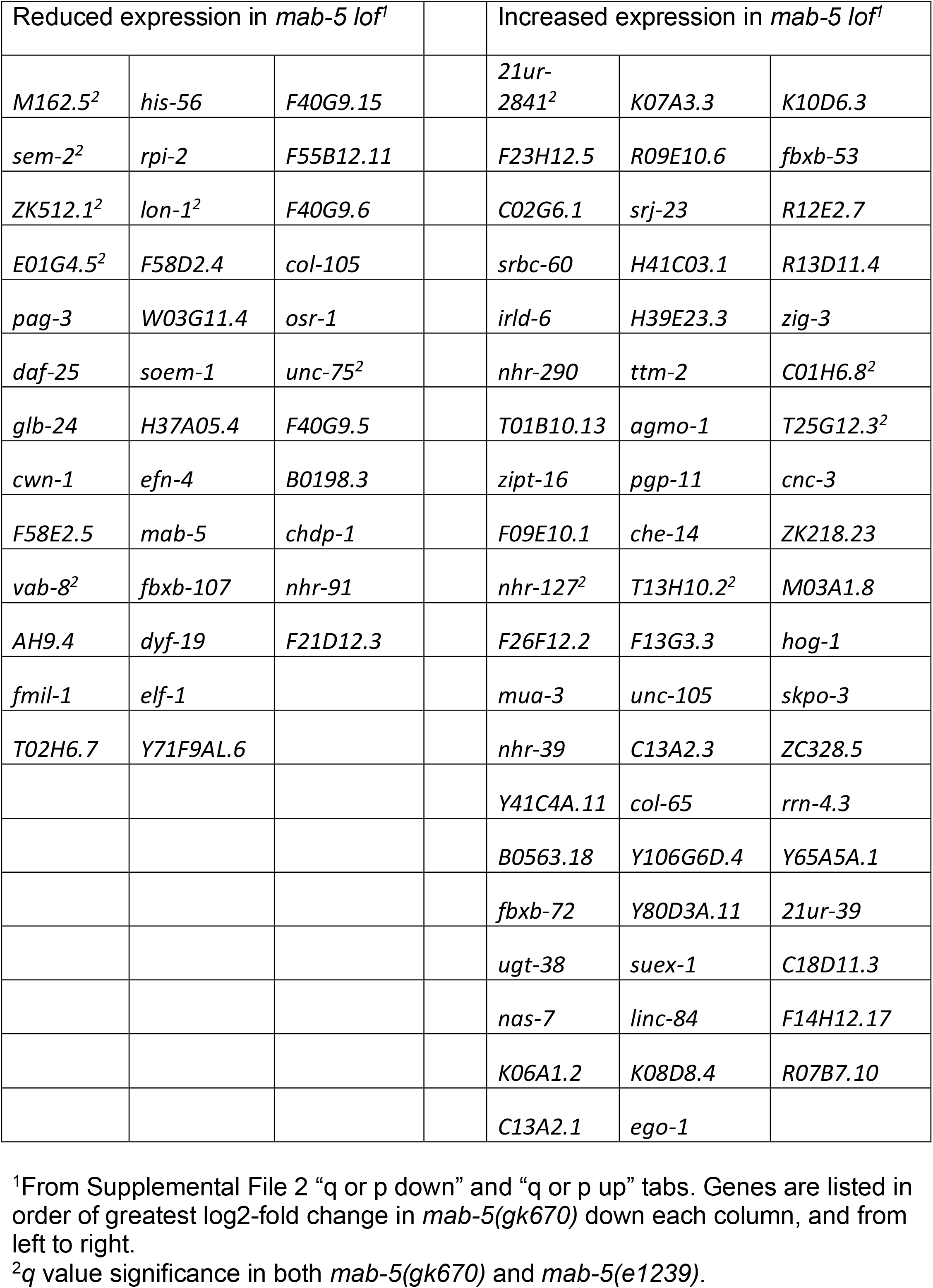
Genes affected in Q cells from *mab-5* loss-of-function.

Combined, 238 genes showed significantly reduced expression in one or both mutants, and 116 showed increased expression in one or both (Supplemental File 2; Down in either and Up in either tabs). Far fewer genes overall were identified in the *mab-5(e1239)* analysis. Possibly, *mab-5(e1239)* is hypomorphic and has a less severe effect on gene expression compared to *mab-5(gk670)*. However, the PQR migration phenotype is similar in the two alleles (Table 1). *mab-5* itself showed significantly-reduced expression in *mab-5(gk670)*. In *mab-5(e1239)*, *mab-5* expression was reduced, but with only *p-*value significance and not *q-*value significance, consistent with *mab-5(e1239)* being a potential hypomorph.

To increase stringency and to narrow the gene list, genes with *q* value significance in one *mab-5* mutant and either *q* or *p* value significance in the other mutant were identified (Supplemental File 2; “q or p down” and “q or p up” tabs). In this analysis, 37 genes were downregulated and 59 genes upregulated (Table 2). MAB-5 is known to inhibit the expression of *lin-39* and *mig-13* in QL and descendants (SALSER *et al*. 1993; WANG *et al*. 2013). Neither *lin-39* nor *mig-13* expression in Q neuroblasts was significantly affected in either *mab-5(gk670)* or *mab-5(e1239).* This could reflect that the resolution of our analysis is insufficient to detect expression in one versus both Q cells.

The Database for Annotation, Visualization, and Discovery (DAVID) (HUANG DA *et al*. 2009) was used to analyze the 37 genes downregulated in Table 2 using the Functional Annotation Clustering tool. Among downregulated genes, five significantly-enriched functional clusters were identified (Supplemental File 3 and Figure 4), including terms involved in neurogenesis, transcriptional regulation, and plasma membrane proteins. Among the 59 genes upregulated in Table 2 in *mab-5(lof)*, four functional clusters were identified (Supplemental File 3 and Figure 4), including glycosyltransferase activity and nuclear hormone receptor signaling.

**Figure 4.**
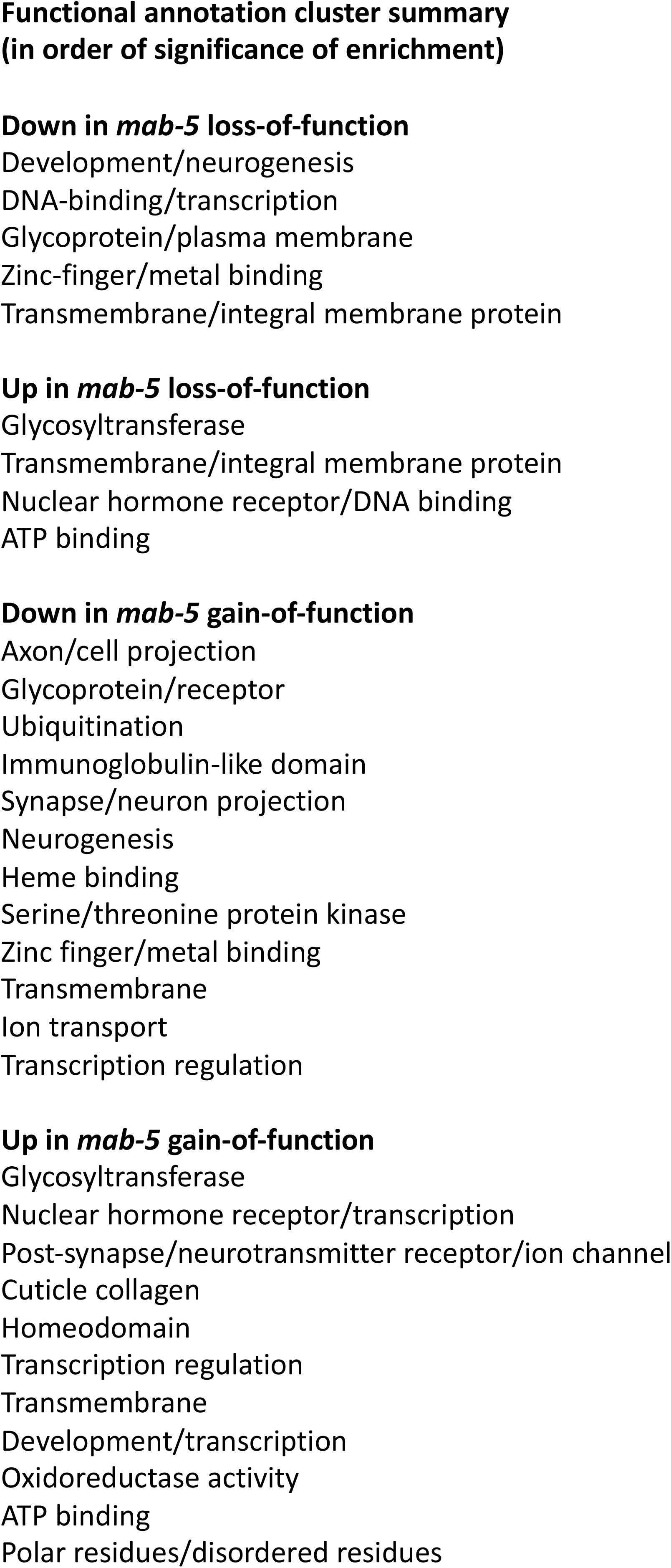
A summary of functional annotation clusters among genes differentially expressed in *mab-5* loss-of-function and gain-of-function mutants. A one-word summary for each of the functional annotation clusters in Supplemental File 3 are listed (genes with expression differences with q value significance in one *mab-5* mutant and at least *p* value significance in the other).

### Differentially expressed genes in *mab-5* gain-of-function mutant Q cells

Two strains with *mab-5(gof)*were analyzed, *lqIs221* and *mab-5(e1751)*. *lqIs221* is a transgene that expresses *mab-5* in both QL and QR under the control of the *egl-17* promoter (JOSEPHSON *et al*. 2016a). *mab-5(e1751)* is a chromosomal gain-of-function mutation (SALSER *et al*. 1993; JOSEPHSON *et al*. 2016a). Genomic DNA from *mab-5(e1751)* was sequenced, aligned to the reference genome, and analyzed using the Integrated Genome Viewer (see Materials and Methods). A 170,672-bp region of chromosome III containing *mab-5* was duplicated in *mab-5(e1751)* (Figure 5), consistent with *mab-5(e1751)* being a large tandem duplication of *mab-5* and other genes (SALSER *et al*. 1993). The duplicated region starts 3,588 bp upstream of the *mab-* 5 start codon between *mab-5* and the downstream gene *C08C3.*4a). In both *mab-5(gof)*, 90-92% of AQR migrate posteriorly to the normal position of PQR, and the remainder fail to migrate from the QR birthplace (Table 1).

**Figure 5.**
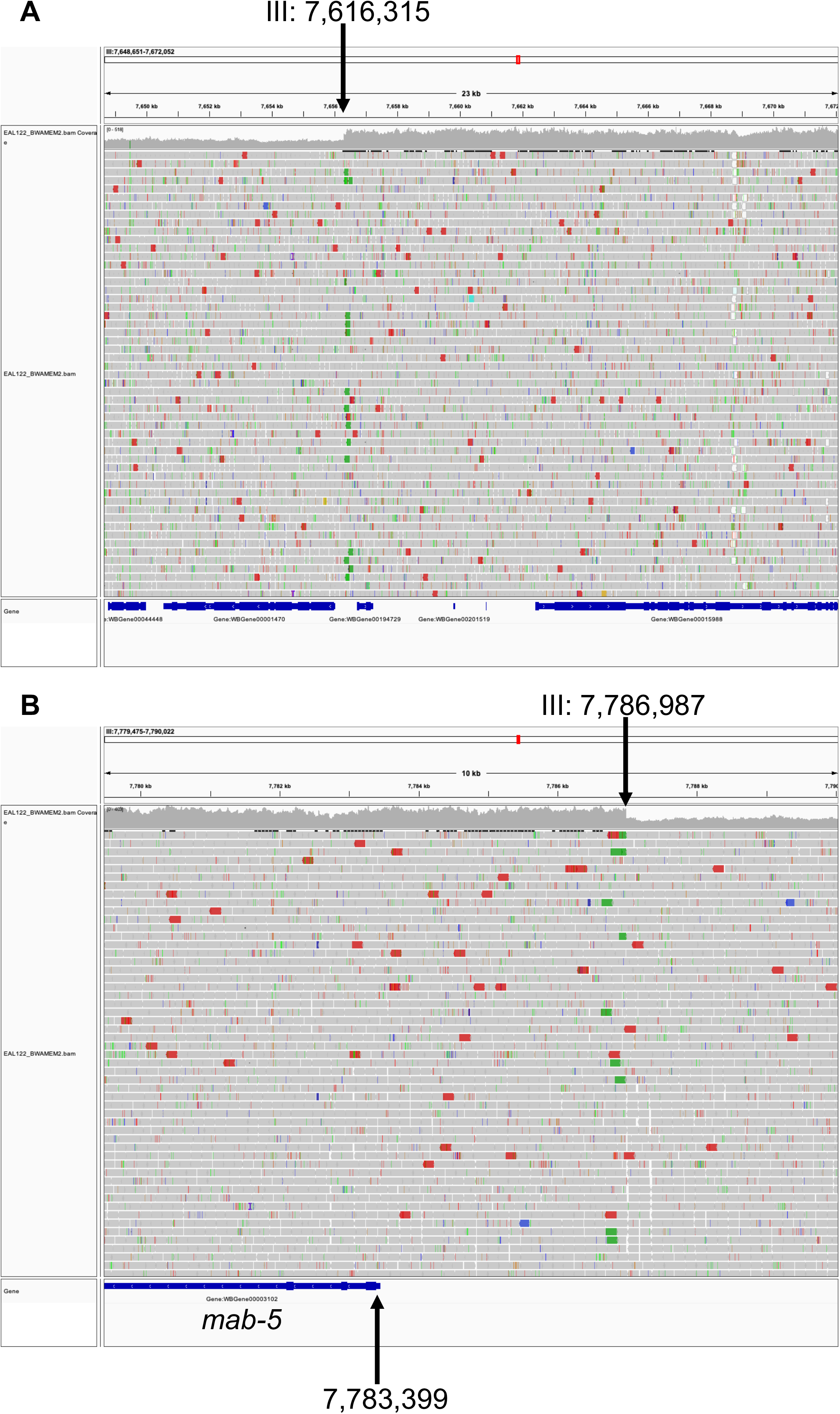
The *mab-5(e1751)* tandem duplication. Genomic DNA from *mab-5(e1751)* was sequenced using Illumina paired-end sequencing to an average read depth of ∼125. The Integrated Genome Viewer was used to analyzed the aligned reads. A 170,672-bp region of chromosome III containing *mab-5* was duplicated in *mab-5(e1751)* (Figure 6) (average read depth of ∼350). A) The 5’ end of the duplicated region with start of the duplication indicated. B) The 3’ end of the duplication with the end site indicated. The *mab-5* gene is also indicated.

In *lqIs221* Q cells, 157 genes had significantly reduced expression (*q*-value ≤ 0.05, log2-fold change ≥ 0.58, base mean expression ≥ 10), and 152 had increased expression (Supplemental File 4). In *mab-5(e1751)*, 121 genes showed reduced Q cell expression, and 276 showed increased Q cell expression (Supplemental File 4). When *q* value significance in one mutant and at least *p* value significance in the other mutant is considered, 91 genes were downregulated in *mab-5(gof)* and 178 upregulated (Supplemental File 4 and Table 3). A majority of genes showed *q* value significant changes in both mutants (Supplemental file 4). *mab-5* expression was increased in both *e1751* (*p-*value significance) and *lqIs221*(*q-*value significance), consistent with both mutant conditions causing an increase in *mab-5* expression (Supplemental File 4 and Table 3)*. lin-39* expression was reduced in both *e1751* (*p*-value) and *lqIs221* (*q*-value). *mig-13* expression was reduced with *q*-value significance in both. These results are consistent with previous results showing that MAB-5 inhibits expression of *lin-39* and *mig-13* in the Q cells (SALSER *et al*. 1993; WANG *et al*. 2013).

**Table 3.**
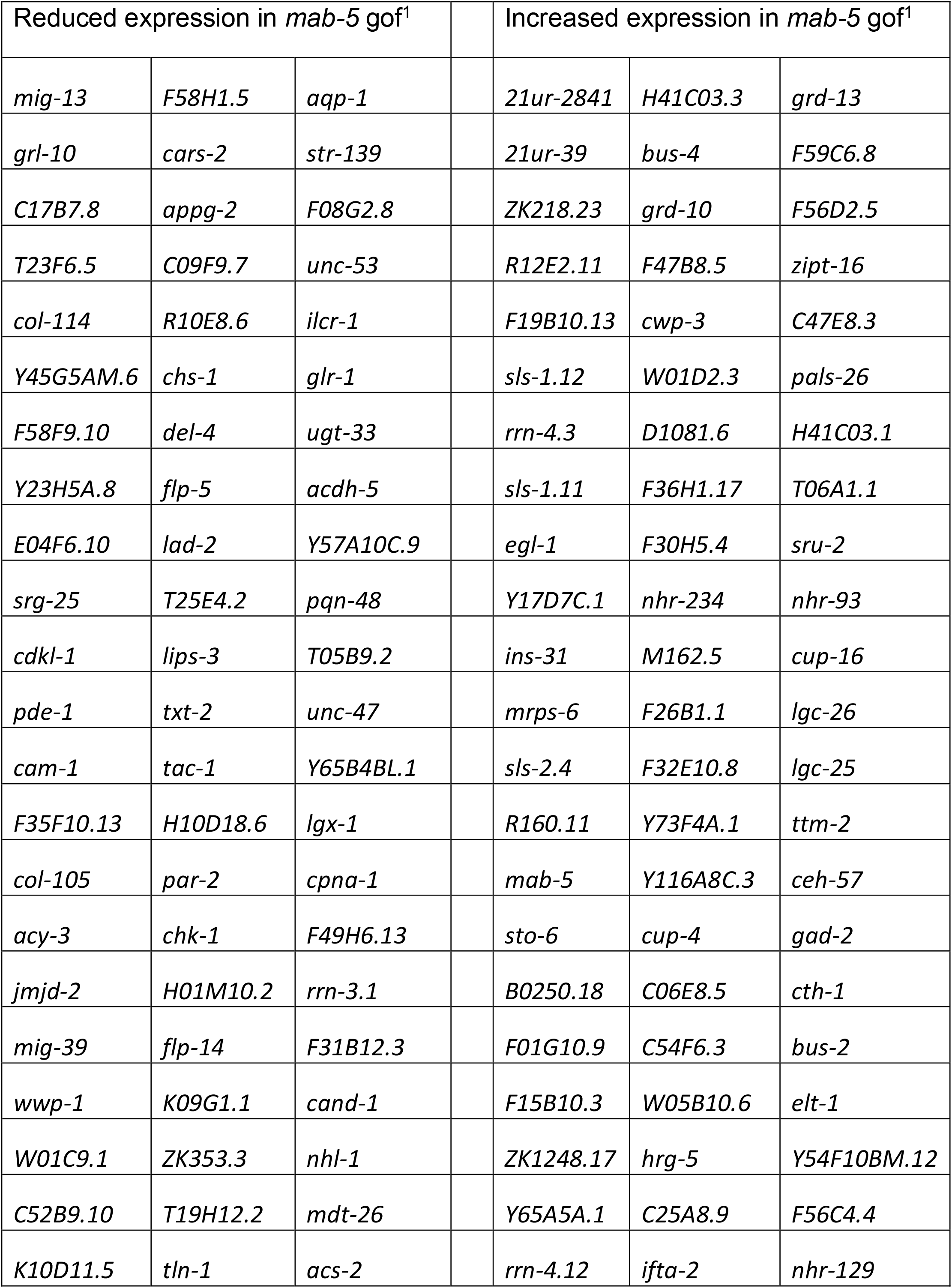
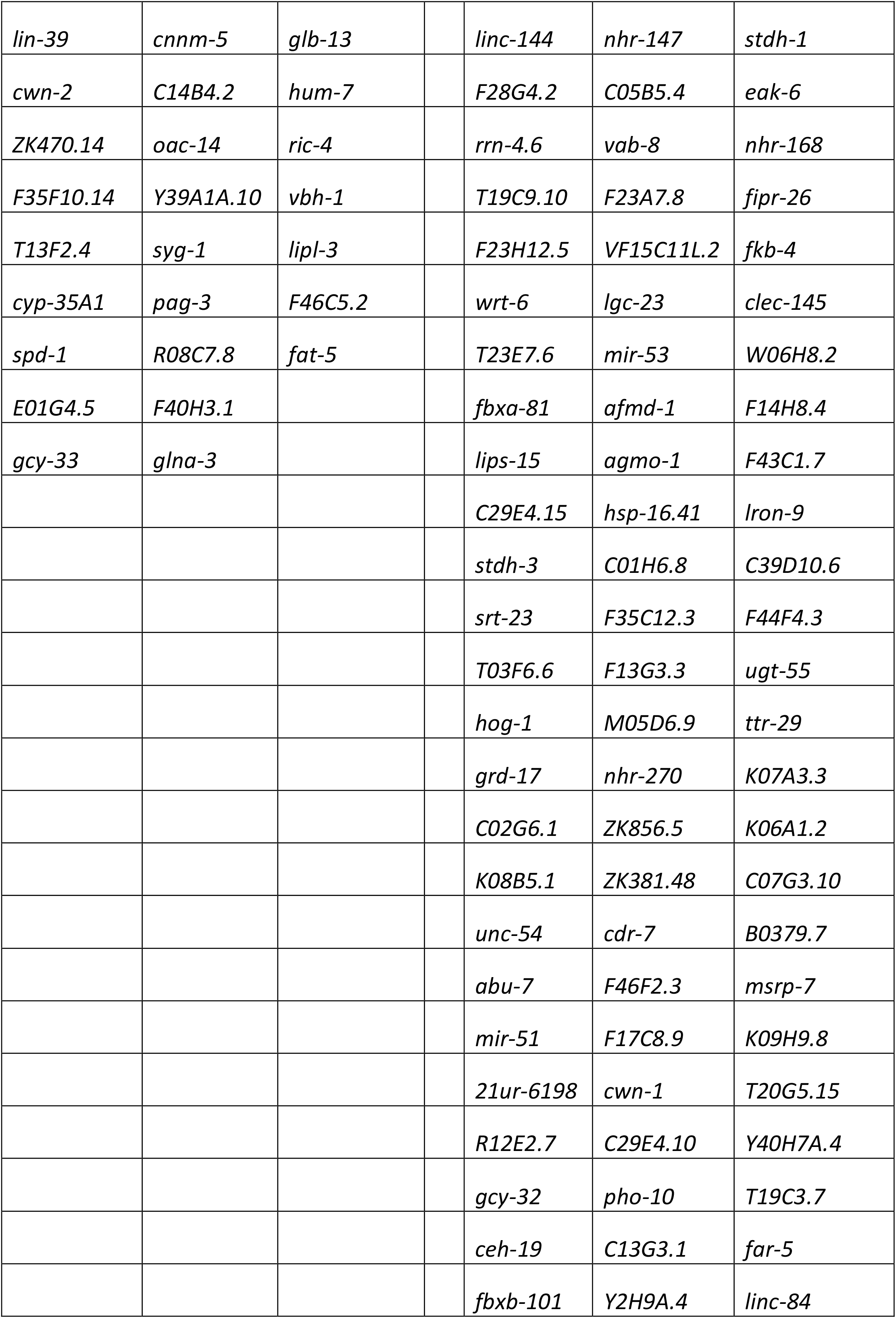
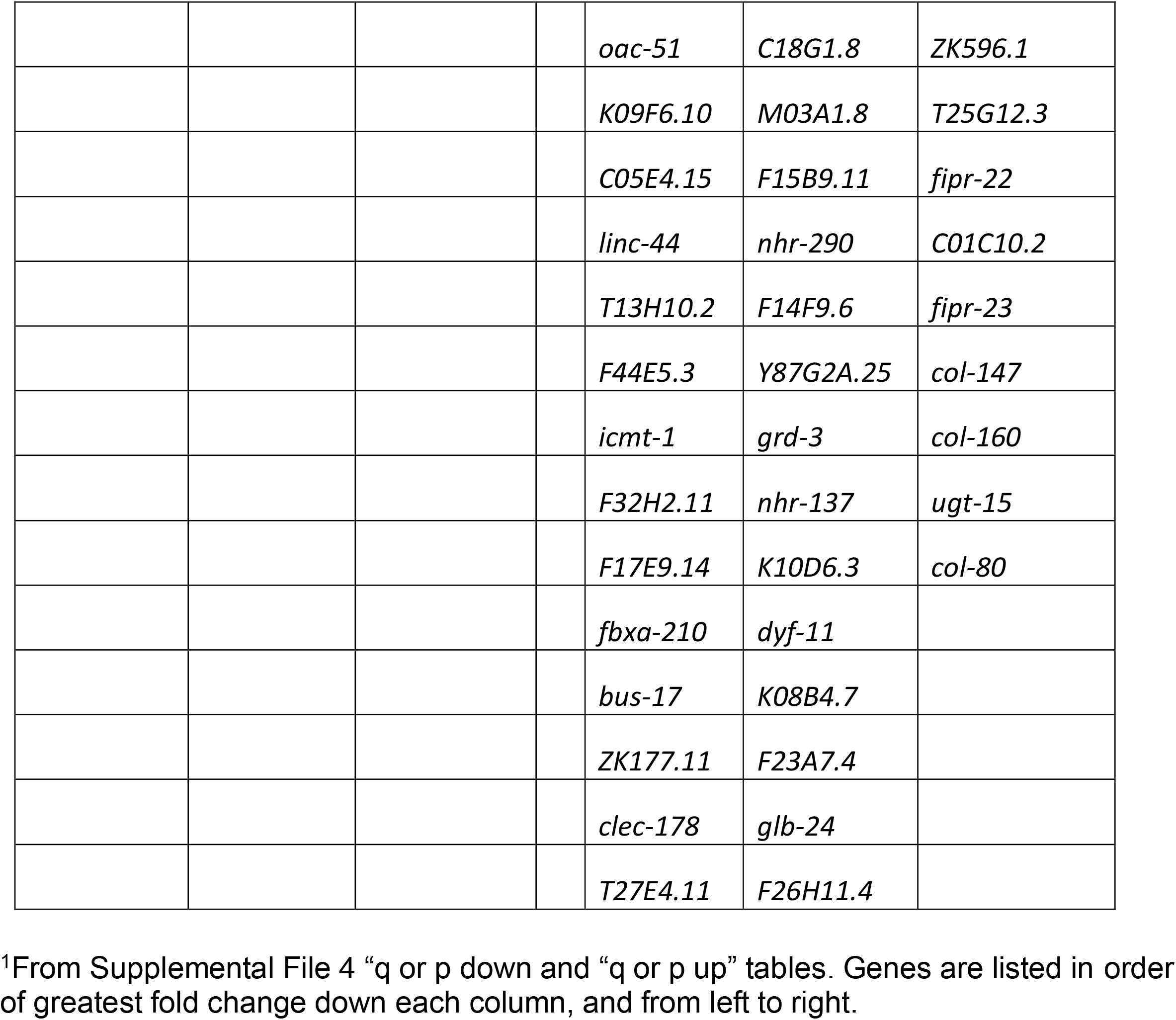
Genes affected in Q cells from *mab-5* gain-of-function.

Functional analysis clustering of the 91 genes downregulated in *mab-5(gof)* identified 12 significant functional clusters, including neurogenesis and neuronal function, glycoprotein, transmembrane protein, and kinase function (Supplemental File 3 and Figure 4). The 178 genes upregulated in *mab-5(gof)* clustered in 11 functional groups, including glycosyltransferase, transcription, neuronal function, and collagen (Supplemental File 3 and Figure 4).

### Paired expression responses in *mab-5* loss-of-function and gain-of-function

Genes with paired responses (i.e., up in *mab-5(lof)* and down in *mab-5(gof)*, and *vice versa*) were identified. Eleven genes that were reduced in one or both *mab-5(lof)* mutants were increased in one or both *mab-5(gof)* mutants, and two genes that were up in *mab-5(lof)* were reduced in *mab-5(gof)* (Table 4). This relatively small number of genes with paired responses is similar to what was observed with *mab-5* RNA-seq from whole animals (TAMAYO *et al*. 2013), and suggests that gene regulation involving MAB-5 is complex and involves other factors in addition to MAB-5.

**Table 4.** Paired responses to *mab-5*.

| Down in lof, up in gof |  |  |  | Up in lof, down in gof |  |
| --- | --- | --- | --- | --- | --- |
| *F59B10.2* | *cwn-1* | *ZK682.7* |  | *mua-3* | *str-139* |
| *dyf-5* | *vab-8* | *ifta-2* |  |  |  |
| *M162.5* | *gcy-32* | *ZK512.1* |  |  |  |
| *mab-5* | *glb-24* |  |  |  |  |

### Genes involved in initial Q cell polarization and migration were not differentially expressed

Genes that have been shown to function in initial QR/QL asymmetric polarization and migration were not differentially expressed in our *mab-5* mutant analyses. This includes the transmembrane proteins UNC-40/Deleted in Colorectal Cancer (DCC), DPY-19/mDpy19, MIG-21, MIG-15/NIK, PTP-3/LAR, and CDH-4 [reviewed in (RELLA *et al*. 2016)]. This was an expected result, as these genes affect both QL and QR migration, and initial polarization and migration is MAB-5-independent.

The heparan sulfate proteoglycan (HSPG) LON-2 and an enzyme that modifies HSPGs, HSE-5, have also been shown to function in early Q neuroblast protrusion and migration (SUNDARARAJAN *et al*. 2015; WANG *et al*. 2015). Neither of these genes were significantly differentially expressed in our *mab-5* mutant analyses. Genes involved in cytoskeleton remodeling and cell adhesion expected to act in all cell migration events were also not found to be differentially expressed in our analyses, including: *ced-10/Rac, mig-2/RhoG, unc-73/Trio, pix-1/PIX, unc-34/Ena, ani-1/Anillin, cor-1/Coronin,* and *ina-1/Integrin⍺*.

### Wnt receptors and cytoplasmic signaling molecules are expressed in Q cells

Relative Wnt pathway gene expression (Table 5, adapted from SAWA AND KORSWAGEN (2013)) in Q cells was determined by base mean expression in Supplemental File 1 “WTQ x WTall” tab. Genes involved in Wnt secretion and Wnt ligand genes were expressed at relatively low levels, where Wnt receptors and cytoplasmic and nuclear signaling components were expressed at relatively high levels. The Wnt receptors *mom-5, cfz-2*, and *lin-18* were expressed relatively lower than *lin-17, mig-1,* and *cam-1*. *bar-1* and *pop-1* expression were relatively low, surprising given their centrality to Wnt signaling and *mab-5* expression in the Q neuroblasts. The secreted Wnt inhibitor *sfrp-1* and components of planar cell polarity were also highly expressed in the Q neuroblasts.

**Table 5.** Effects of *mab-5* on expression of Wnt signaling genes in the Q cells.

| **Component** | ***C. elegans* ortholog** | **Base mean expression<sup>1</sup>** | **RNA-seq Results** |
| --- | --- | --- | --- |
|  | **Wnt production and secretion** |  |  |
| Porcupine | *mom-1* | 185 | NO |
| Wntless | *mig-14* | 644 | NO |
| Retromer complex | *vps-26, vps-29, vps-35, snx-3* | 644,<br>428,<br>1080,<br>60 | NO |
|  | **Wnt ligands** |  |  |
|  | *mom-2* | 56 | NO |
|  | *lin-44* | 651 | NO |
|  | *egl-20* | 138 | NO |
|  | *cwn-1* | 490 | UP in GOF<br>DOWN in LOF |
|  | *cwn-2* | 859 | DOWN in GOF |
|  | **Wnt receptors** |  |  |
| Frizzled | *mom-5* | 393 | NO |
|  | *lin-17* | 1982 | DOWN in LOF |
|  | *mig-1* | 1801 | DOWN in GOF |
|  | *cfz-2* | 518 | NO |
| Ror | *cam-1* | 4489 | DOWN in GOF |
| Ryk/Derailed | *lin-18* | 282 | DOWN in LOF |
|  | **Secreted Wnt inhibitors** |  |  |
| Sfrp | *sfrp-1* | 1144 | NO |
|  | **Wnt/β-catenin pathway components** |  |  |
| CK1 | *kin-19* | 6726 | NO |
| Dishevelled | *mig-5* | 859 | NO |
|  | *dsh-1* | 2363 | NO |
|  | *dsh-2* | 3459 | NO |
| GSK3β | *gsk-3* | 1212 | NO |
| Axin | *pry-1* | 2769 | NO |
|  | *axl-1* | 238 | NO |
| APC | *apr-1* | 1486 | NO |
| βTrcp | *lin-23* | 3379 | NO |
| Nlk | *lit-1* | 25730 | NO |
| Tak1 | *mom-4* | 5714 | NO |
| Tab1 | *tap-1* | 2286 | NO |
| β-catenin | *bar-1* | 667 | DOWN in GOF |
|  | *sys-1* | 2990 | NO |
|  | *wrm-1* | 3077 | NO |
|  | *hmp-2* | 3892 | NO |
| Tcf | *pop-1* | 272 | DOWN in LOF |
| Groucho | *unc-37* | 3187 | NO |
|  | **PCP pathway components** |  |  |
| Vangl | *vang-1* | 391 | NO |
| Prkl | *prkl-1* | 1402 | NO |
| Flamingo | *fmi-1* | 37512 | NO |

<sup>1</sup>Rounded to the whole number from Supplemental File 1 “WTQ vs WTall” tab.

### MAB-5 regulates Wnt ligand and receptor expression

In QL, EGL-20/Wnt is required to activate MAB-5 expression. In QR, anterior migration of the QR descendants is dependent on non-canonical, β-catenin-independent signaling pathways, which do not result in the expression and activation of MAB-5 (ZINOVYEVA AND FORRESTER 2005; MENTINK *et al*. 2014; SCHILD *et al*. 2023). Furthermore, MAB-5 participates in feedback regulation to adjust Wnt-signaling responses (JI *et al*. 2013). The effects on *mab-5* on Wnt pathway genes was analyzed (Table 5, adapted from SAWA AND KORSWAGEN (2013). If an effect was observed in one or both mab-5*(lof)* or *mab-5(gof)* mutants, it was noted on Table 5.

*cwn-1/Wnt* showed a paired response, with decreased expression in *mab-5(lof)* and increased expression in *mab-5(gof)*. *cwn-1* mutants display failure of AQR to migrate fully (Table 1), but no effects on PQR alone. *cwn-2* showed decreased expression *mab-5(gof)*, but alone had no effect on AQR or PQR migration, but might have redundant roles with other *Wnt* genes in AQR and PQR migration (JOSEPHSON *et al*. 2016a).

There are four Frizzled receptors in the canonical Wnt/β-catenin pathway: MIG-1, LIN-17, CFZ-2, and MOM-5. Of these Frizzled receptors, MIG-1, LIN-17, and MOM-5 have been previously shown to be required for *mab-5* expression in QL (HARRIS *et al*. 1996). *mig-1* expression was decreased in *mab-5(gof)*. This suggests that MAB-5 represses expression of *mig-1*. This is consistent with previous data as MIG-1 has been shown to be a target of MAB-5 (NIU *et al*. 2011). Furthermore, evidence of a negative feedback loop between MIG-1 and the Wnt signaling pathway has been previously shown (SCHILD *et al*. 2023), although the negative transcriptional regulation of MIG-1 was proposed to be MAB-5 independent (JI *et al*. 2013). However, based on our data, it is possible that MAB-5 is directly involved in regulating MIG-1 transcription via a negative feedback loop. *lin-17* was also decreased in *mab-5(lof)*.

Two additional receptors involved in non-canonical Wnt signaling include Ror (CAM-1) and Ryk/Derailed (LIN-18). *cam-1* displayed decreased expression in the *mab-5(gof)* (Table 5), indicating that MAB-5 inhibits expression of *cam-1.* Previous work showed that in *cam-1(gm122)* mutants, QR descendants (QR.pax) displayed weak undermigration, while QL descendant migration appeared unaffected (KIM AND FORRESTER 2003; MENTINK *et al*. 2014). Previous work also found that overexpression of wild-type *cam-1* or of any *cam-1* derivative that still contained its cysteine-rich domain caused QL descendants to migrate anteriorly (KIM AND FORRESTER 2003), and overexpression of *cam-1* resulted in a decrease in *mab-5* expression in QL (FORRESTER *et al*. 2004). Taken together, these data suggest that MAB-5 might reciprocally inhibit expression of *cam-1* and that CAM-1 function is needed to promote anterior migration of the Q cell descendants. *lin-18* displayed decreased expression in *mab-5(lof)*, but *lin-18* mutants showed no AQR or PQR migration defects (0%, n = 100).

Other key components of canonical Wnt/β-catenin signaling are listed in Table 5, including canonical cytoplasmic signaling molecules and planar cell polarity. Of these, *bar-1/β-catenin* was down in *mab-5(gof)* and *pop-1/TCF* was down in *mab-5(lof)*. While both *bar-1* and *pop-1* are required for PQR posterior migration, the functional significance of these expression differences is unclear. In sum, MAB-5 might modulate Wnt signaling in the QL lineage predominantly by influencing expression of Wnt ligands and receptors.

### *cwn-1/Wnt* inhibits anterior QL.a/p migration with *egl-20/Wnt*

*cwn-1/Wnt* expression in Q cells showed a strong paired response to *mab-5* mutation. *cwn-1* expression was reduced in *mab-5(lof)* and increased in *mab-5(gof)* (Table 4 and 5) suggesting that MAB-5 is necessary and sufficient for *cwn-1* expression in Q cells. *cwn-1(ok546)* mutants displayed failure of AQR to migrate fully, but showed no PQR migration defects indicative of acting with *mab-5* (Table 1). Two other predicted null alleles of *cwn-1* showed a similar phenotype (Table 1).

Given the redundant function of *Wnt* genes in *C. elegans* (ZINOVYEVA AND FORRESTER 2005; JOSEPHSON *et al*. 2016a), the role of *cwn-1* in PQR migration might be masked. *egl-20* null alleles result in a near-complete migration of PQR anteriorly to the normal position of AQR (*e.g., egl-20(lq42)* in Table 1). However, *egl-20(mu39)* is a hypomorphic missense mutation that results in 48% of PQRs migrating normally to position 5 in the tail (Table 1). *egl-20(mu39); cwn-1* double mutants displayed significantly more severe PQR anterior migration defects than *egl-20(mu39)* alone (1-3% position 5 compared to 48%), indicating that *cwn-1* enhanced the PQR migration defects of hypomorphic *egl-20(mu39)*. This suggests that CWN-1 might have a role along with EGL-20 to inhibit anterior QL.a/p migration. Misdirected PQRs often failed in anterior migration similar to AQR, suggesting that the misdirected PQRs are subject to the same effects of *cwn-1* mutation on anterior migration as are AQRs.

### *cwn-1* acts downstream of MAB-5

The transcriptomic analysis presented here suggests that MAB-5 is necessary and sufficient for *cwn-1* expression in the Q cells and descendants. If *cwn-1* acts downstream of MAB-5, it might suppress the effects of *mab-5(gof)*. In *mab-5(e1751)* and *lqIs221* gof, 90% and 92 of AQR migrate posteriorly to the normal position of PQR (position 5) (Table 1). *lqIs221* and *mab-5(e1751)* double mutants with the predicted null *egl-20(lq42)* showed 11 and 3 AQRs migrating anteriorly, never observed in *lqIs221* or *mab-5(e1751)* alone. *lqIs221* double mutants with *egl-20(gk453010),* also a null, showed anterior AQR migration as well (Table 1). These data suggest that *egl-20* might act in part downstream of *mab-5* to inhibit anterior migration.

*cwn-1(ok546)* alone had no effect on either *mab-5(gof)* condition (Table 1). However, *egl-20(lq42); cwn-1(ok546); mab-5(e1751)* and *egl-20(lq42); cwn-1(ok546); lqIs221)* triple mutants displayed strong anterior AQR migration, with only 9% and 25 % migrating posteriorly Furthermore, anterior PQR migration was also observed (Table 1), never seen in *lqIs221* or *mab-5(e1751)* alone. These results indicate that both *cwn-1* and *egl-20* are required for the ability of *mab-5(gof)* to inhibit anterior migration. These results also suggest that *egl-20* is required after *mab-5* is activated, possibly in parallel to *cwn-1.* Combined with the Q cell transcriptomic data, these results indicate that CWN-1 normally acts downstream of MAB-5 in QL.a/p to inhibit anterior QA.a/p migration, in parallel to EGL-20 (Figure 6).

**Figure 6.**
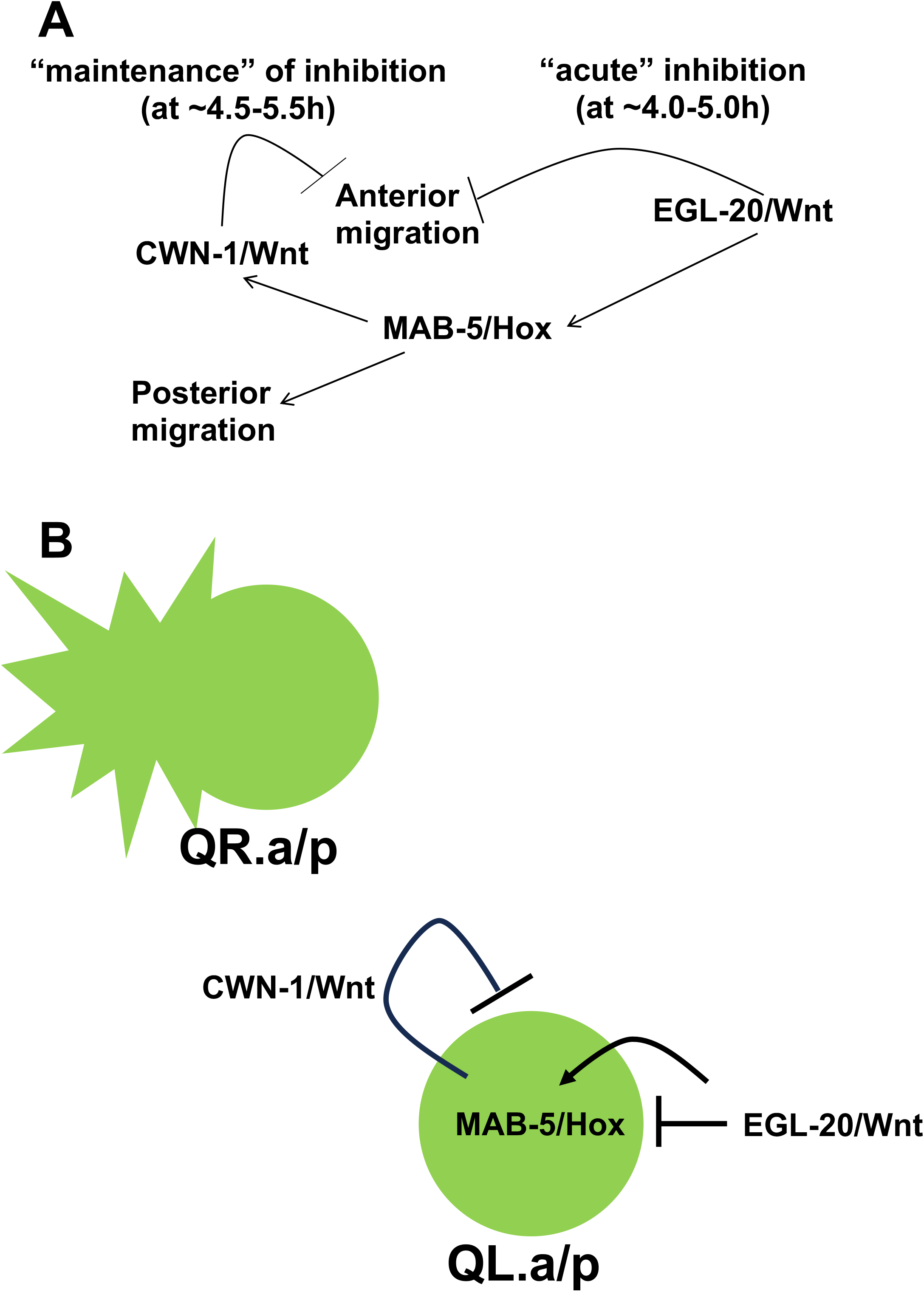
A model of the role of MAB-5 and CWN-1 in QL.a/p. A) A pathway diagram of MAB-5 function. B) A cellular diagram of MAB-5 function. In the absence of the EGL-20/Wnt signal, QR.a/p polarizes and migrates anteriorly, as do QL.a/p from *egl-20* and *mab-5* mutants. EGL-20/Wnt acutely inhibits anterior migration of QL.a/p via a non-canonical Wnt signaling pathway, and activates MAB-5 expression in QL.a/p via canonical Wnt signaling. *cwn-1* expression in QL.a/p is activated by MAB-5, and CWN-1 acts in an autocrine manner to maintain inhibition of anterior migration, in parallel to EGL-20.

## Discussion

Neuroblast and neuronal differentiation rely on the precise and discrete actions of many genes that act together to coordinate development. In *C. elegans*, the MAB-5/Hox transcription factor controls the direction of Q neuroblast migration without obviously affecting other aspects of differentiation. In an effort to understand the role of a terminal selector Hox gene on neuroblast migration, the transcriptomes of Q cells from wild-type and *mab-5* loss- and gain-of-function mutants were determined using fluorescence-activated cell sorting and RNA-seq.

### Transcriptome of a differentiating neuroblast lineage

Q cells in the early stages of their differentiation were isolated and subject to transcriptomic analysis. The populations that were sequenced included both QL and QR cells in different phases of early development, as well as their anterior and posterior daughters after the first cell division (Figure 1). This is well before the final differentiation of any lineage descendant into neurons, and before the final migrations of the neuroblasts and neurons. Presence of genes known to be expressed in Q neuroblasts indicates that they were successfully purified and sequenced (Figure 3). However, significant temporal and cell-type heterogeneity in the experiment likely missed finer details of discrete steps of Q differentiation and migration.

### Genes regulated by MAB-5/Hox in the Q cells

Genes whose expression was affected by *mab-5(lof)* and *mab-5(gof)* were identified using two independent lof and gof mutant backgrounds. Common genes were found, but there was also much heterogeneity in the genes in the different backgrounds. This could reflect difference in mutant effect (*e.g., mab-5(e1239)* might be hypomorphic compared to *mab-5(gk670)*). It could also be due to experimental variability between different strains.

The functional annotation profile of genes regulated by MAB-5 are generally those expected of a differentiating neuroblast lineage. While this analysis reveals genes that are regulated by *mab-5*, it is likely that many genes have been missed in this analysis due to experimental variability and heterogeneity of cell type and time. For example, it is known that *mab-5* inhibits *lin-39* in QL. However, *lin-39* expression was not significantly increased in *mab-5(lof)*, although was decreased in *mab-5(gof)*. Thus, genes from both *mab-5(lof)*and *mab-5(gof)*analyses should be considered as targets of MAB-5 despite not showing a paired response. Indeed, very few genes showed a paired response in *mab-5(lof)*versus *mab-5(gof)*conditions. In addition to experimental variability, this could also be due to different interactions with other factors required for expression of those genes.

### MAB-5 targets involved in posterior migration

A goal of these studies was to identify genes that act downstream of MAB-5 to drive posterior Q lineage migrations. Previous studies showed that the effect of MAB-5 on posterior migration is two-fold. First, MAB-5 is required to prevent anterior migration. Second, MAB-5 is required to reprogram QL.a to migrate posteriorly. MAB-5 could stimulate expression of genes to accomplish these feats, and/or could repress expression of genes required for anterior migration. *lin-39* and *mig-13* are required for anterior migration and are known repressive targets of MAB-5 in QL. Here, expression of *lin-39* and *mig-13* were reduced *mab-5(gof)*, consistent with MAB-5 repressing genes required for anterior migration.

### CWN-1/Wnt is a MAB-5 target that inhibits anterior migration

Genes down in *mab-5(lof)* and/or up in *mab-5(gof)* could include genes that are activated by MAB-5 to inhibit anterior migration or promote posterior migration. *cwn-1/Wnt* showed a strong paired expression response to *mab-5*, and was down in *mab-5(lof)*and up in *mab-5(gof)*. While *cwn-1* mutants alone had no effect on PQR migration, they were strong enhancers PQR anterior migration of a hypomorphic *egl-20* mutant. Furthermore, *egl-20; cwn-1* double mutants suppressed posterior AQR (and PQR) migration in *mab-5(gof)* mutants, suggesting that CWN-1/Wnt acts downstream of MAB-5 and is required for MAB-5 to inhibit anterior migration.

A revised model of MAB-5 and CWN-1 function is shown in Figure 6A (JOSEPHSON *et al*. 2016a). Previous studies showed that EGL-20 has a dual role in posterior QL migration. Via non-canonical Wnt signaling, EGL-20 acutely inhibits anterior migration of QL.a/p. Via canonical Wnt signaling, EGL-10 drives expression of MAB-5. In turn, MAB-5 maintains the inhibition of anterior migration, and also reprograms QL.a to migrate to the posterior. Work presented here indicates that MAB-5 drives expression of CWN-1 in the Q cells to inhibit anterior migration, likely in parallel to EGL-20 acute inhibition. Thus, CWN-1 might act in an autocrine manner in the Q cells to inhibit anterior migration, possibly via a non-canonical Wnt pathway (Figure 6B). The nature of this non-canonical pathway, as well as the nature of genes that promote posterior migration downstream of MAB-5, might be revealed through further analysis of MAB-5 target genes defined here. These studies on *cwn-1* are proof-of-principle that the gene lists generated in this transcriptomic analysis of Q cells in *mab-5* mutants contains other genes that act downstream of MAB-5 to control posterior migration.

## Supporting information

Q cell gene expression

Genes affected by mab-5 lof

Functional annotation clustering

Genes affected by mab-5 gof

## Acknowledgments

The authors thank members of the Lundquist and Ackley labs for discussion, and the CGC, which is funded by NIH Office of Research Infrastructure Programs (P40 OD010440) for some strains. This work was supported by NIH grant R01 NS115467 to E.A.L., and the NIH Kansas Infrastructure Network of Biomedical Research Excellence (P20 GM103418). Whole genome sequencing was conducted at the KU Genome Sequencing Core, which is part of the NIH *Center for Molecular Analysis of Disease Pathways* (P30 GM145499).

